# Optimization of multiplexed CRISPR/Cas9 system for highly efficient genome editing in *Setaria viridis*

**DOI:** 10.1101/2020.04.11.037572

**Authors:** Trevor Weiss, Chunfang Wang, Xiaojun Kang, Hui Zhao, Maria Elena Gamo, Colby G. Starker, Peter A. Crisp, Peng Zhou, Nathan M. Springer, Daniel F. Voytas, Feng Zhang

## Abstract

In recent years, *Setaria viridis* has been developed as a model plant to better understand the C4 photosynthetic pathway in major crops. With the increasing availability of genomic resources for *S. viridis* research, highly efficient genome editing technologies are needed to create genetic variation resources for functional genomics. Here, we developed a protoplast assay to rapidly optimize the multiplexed CRISPR/Cas9 system in *S. viridis.* Targeted mutagenesis efficiency was further improved by an average of 1.4-fold with the exonuclease, *Trex2.* Distinctive mutation profiles were found in the Cas9_Trex2 samples with 94% of deletions larger than 10bp, and less than 1% of mutations being insertions. Further analyses indicated that 52.2% of deletions induced by Cas9_Trex2, as opposed to 3.5% by Cas9 alone, were repaired through microhomology-mediated end joining (MMEJ) rather than the canonical NHEJ DNA repair pathway. Combined with the robust agrobacterium-mediated transformation method with more than 90% efficiency, the multiplex CRISPR/Cas9_Trex2 system was demonstrated to induce targeted mutations in two tightly linked genes, *svDrm1a* and *svDrm1b,* at the frequency ranging from 73% to 100% in T0 plants. These mutations were transmitted to at least 60% of the transgene-free T1 plants with 33% of them containing bi-allelic or homozygous mutations in both genes. This highly efficient multiplex CRISPR/Cas9_Trex2 system makes it possible to create a large mutant resource for *S. viridis* in a rapid and high throughput manner, and has the potential to be widely applicable in achieving more predictable MMEJ-mediated mutations in many plant species.

## Introduction

*Setaria viridis* (green foxtail) is an annual diploid C4 panicoid grass with a small genome and the wild relative to *Setaria italica* (foxtail millet), an agriculturally important crop in parts of Africa and Asia (Lata, Gupta and Prasad, 2013). Although historically regarded as an invasive weed, *S. viridis* has recently been developed as an emerging monocot model species to study bioenergy feedstocks and panicoid food crops, such as maize, sorghum, sugarcane and switchgrass, and to better dissect the cellular and biochemical mechanisms of C4 photosynthesis (Brutnell *et al*., 2010). *S. viridis* has many features that make it an attractive model system including a short life cycle, compact stature, reproduction via self-pollination and the ability to generate a high number of seeds (Defelice, 2002; Brutnell *et al*., 2010). Furthermore, the expanding genetic and genomic resources, including diverse germplasm accessions, chemically induced mutant populations, high quality reference genome of the A10.1 variety and the resequenced genomes from more than 600 wild accessions, make it possible to conduct large-scale gene discovery and functional genomics in *S. viridis* (Bennetzen *et al*., 2012; Zhu, Yang and Shyu, 2017; Huang *et al*., 2019). Lastly, as another key factor for a successful model plant system, an efficient agrobacterium-mediated transformation method has been reported in *S. viridis* indicating it is amenable to genetic engineering techniques (Van Eck, 2018; Huang *et al*., 2019; Nguyen *et al*., 2020).

Genome editing has significant potential to expedite gene discovery and functional genomics. A key characteristic of current genome editing technologies is the use of programmable nucleases, such as Meganucleases, Zinc finger nucleases (ZFNs), Transcriptional activator like effector nucleases (TALENs) or CRISPR/Cas9, to create double-stranded DNA breaks (DSBs, or single-stranded nicks in some applications) at targeted loci. The induced DSBs can be exploited to introduce a variety of genomic modifications, such as deletions, insertions and nucleotide substitutions, by using one of two main DNA repair pathways, end joining or homology-directed repair (HDR). The end joining pathways, including non-homologous end joining (NHEJ) and microhomology-mediated end joining (MMEJ), are mostly used to generate insertions/deletions (Indels) at targeted sites, while the HDR pathway is employed to precisely incorporate desired sequences into targeted loci by copying genetic information from cotransformed donor templates (Chen *et al*., 2019).

In recent years, the CRISPR/Cas9 system has become the reagent of choice to achieve efficient genome editing in many plant and animal species due to its simplicity, robust activity, versatility, and multiplexing capability (Yin, Gao and Qiu, 2017). Using this system, several gene knockout resources have been created in rice and maize (Meng et*al.,* 2017; Liu *et al*., 2020). To generate gene knock-out resources in a plant species, high mutagenesis frequency and multiplexing capability, i.e. targeting multiple loci simultaneously, are key factors. The CRISPR/Cas9 system often requires considerable optimization in vector construction, transgene expression, tissue culture and transformation efficiency when adopted in a new plant species (Yin, Gao and Qiu, 2017). Additional strategies can be employed to further improve mutagenesis efficiency. For example, it has been demonstrated that the use of plants with a deficiency in the NHEJ pathway, such as the *Ku70/Ku80* and *Ligase IV* mutants, could significantly enhance the frequency of targeted mutagenesis (Qi *et al*., 2013). Moreover, the simultaneous expression of exonucleases, such as *Trex2,* with CRISPR/Cas9 has been shown to enhance the frequency of targeted mutagenesis up to 2.5-fold in tomato and barley (Čermák *et al*., 2017). As for improving the multiplexing capability of the CRISPR/Cas9 system, one of the most effective strategies thus far is to achieve multiplex CRISPR guide RNA (gRNA) expression from a single polycistronic cassette. To this end, expression of multiple CRISPR gRNAs is driven by a single promoter with each gRNA separated by ribozyme sites, Csy4 recognition sites, or transfer RNA (tRNA) sequences, which can be processed to release individual mature gRNAs for targeting (Tsai *et al*., 2014; Xie, Minkenberg and Yang, 2015). However, several studies have indicated that these multiplexing systems need to be tested and optimized when used in a new species. The polycistronic cassettes may possess varied processing efficacy in different species, and the Csy4 system may result in cytotoxicity (Minkenberg, Wheatley and Yang, 2017; Shiraki and Kawakami, 2018).

Although one example of the CRISPR/Cas9 mediated gene knockouts has been described in *S. viridis,* a highly efficient, multiplexed gene editing system has yet to be reported (Huang *et al*., 2019). In this study, we developed a protoplast-based transient assay for rapidly testing and optimizing the multiplexed CRISPR/Cas9 system in *S. viridis.* This system was also used to test the strategy of co-expression of the *Tre×2* exonuclease to further improve targeted mutagenesis efficiency in *S. viridis.* Finally, the optimized system was validated in stable transgenic plants to achieve highly efficient and heritable knockouts in two tightly linked *S. viridis* genes. The applications of this highly efficient, multiplexed CRISPR/Cas9_Trex2 system were discussed in creating a large genetic mutant resource for *S. viridis* and achieving unique mutations in plant species.

## Results and Discussion

### Development of multiplexed gene editing using *S. viridis* protoplasts

*We* sought to develop a protoplast-based assay for quickly assessing the CRISPR/Cas9 system in *Setaria viridis* (Figure S1). Protoplasts were isolated from young leaves of 14-day old *S. viridis* seedlings. Transformation efficiency was tested using the green fluorescent protein (GFP) reporter driven by two different promoters, the Cestrum yellow leaf curling virus (*CmYLCV*) promoter and the *Ubiquitin 2* promoter from switchgrass (*PvUbi2*) (Figure S2A). Both constructs produced robust GFP expression in about 80% of protoplasts after 24-hours post transformation (hpt) and at nearly 100% frequency after 48 hpt (Figure S2B).

The *S. viridis* protoplast assay system was then used to test and optimize CRISPR/Cas9 constructs targeting endogenous *S. viridis* genes, the *domains rearranged methylose 1α* (*Drm1a*), *domains rearranged methylose 1b* (*Drm1b*), *male sterile 26* (*Ms26*) and *male sterile 45* (*Ms45*) genes, respectively (Figure 1). The coding sequences of each gene were obtained by BLAST searching the reference genome of *S. viridis* accession A10.1 with the sequences of their maize orthologs, *zmDrm1a, zmDrm1b, zmMs26 and zmMs45* (Table S1). Targeted sequences were additionally verified by Sanger sequencing in *S. viridis* accession ME034v, the plant variety used in this study. CRISPR gRNAs were designed to target the 5’ exons or the conserved domains in each gene using CRISPOR (Haeussler *et al*., 2016). Each target site contains a restriction enzyme site overlapping the CRISPR/Cas9 cut site to facilitate the Cleaved Amplified Polymorphic Sequences (CAPS) assay for subsequent genotyping analysis (Figure 1).

**Figure 1.**
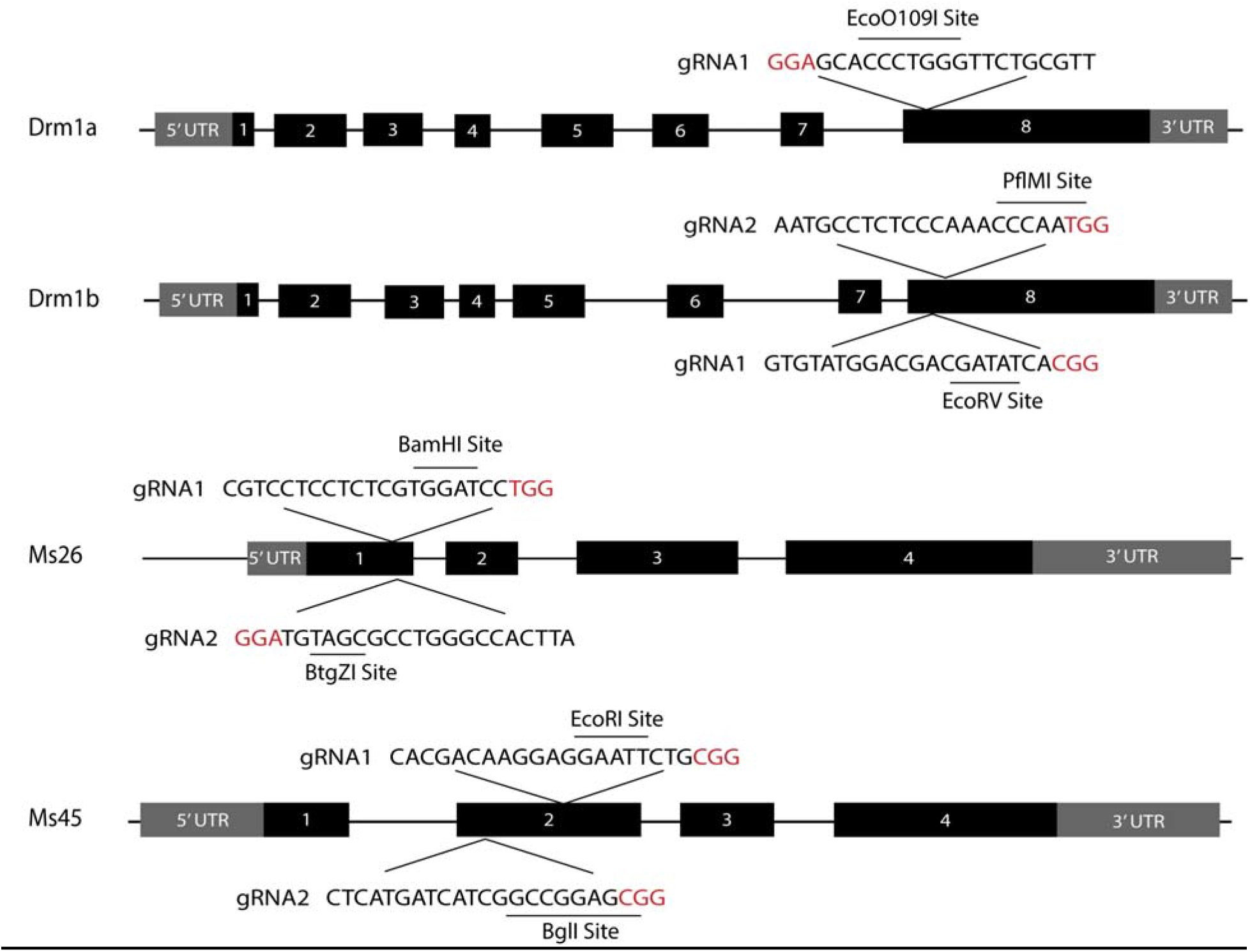
The schematic structures of the *Drm1a, Drm1b, Ms26,* and *Ms45* genes. Each black box represented an exon, with gray boxes representing the 5’ and *3’* untranslated regions. Individual gRNA targeted sites were shown in each gene with the restriction enzyme sites underlined, and the Protospacer Adjacent Motif (PAM) in red.

To achieve multiplexed gene editing in *S. viridis, we* tested two polycistronic gRNA expression systems, the Csy-type (CRISPR system yersinia) ribonuclease 4 (Csy4)-based and the tRNA array-based systems, in protoplasts (Xie, Minkenberg and Yang, 2015; Čermák *et al*., 2017). Constructs containing gRNAs targeting the *Drm1a* and *Drm1b* genes (Figure S3A) were each co-transformed with Cas9 plasmids into protoplasts. As depicted in Figure 2A, high indel mutation frequencies were observed at each target site, ranging from 46% to 82%, indicating that both Csy4 and tRNA-based systems worked effectively in *S. viridis* protoplasts.

**Figure 2.**
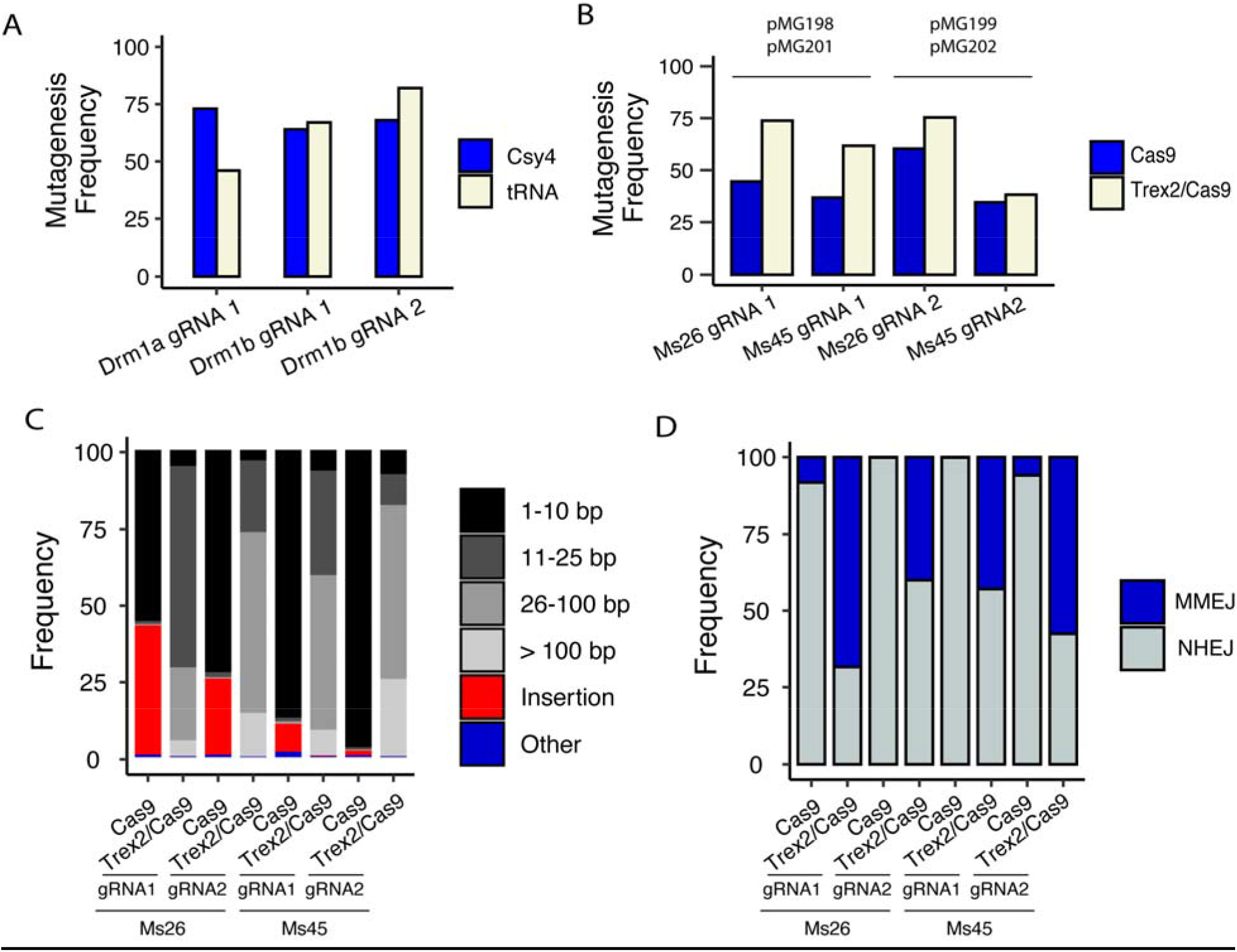
Analyses of mutagenesis frequencies and mutation profiles induced by the CRISPR/Cas9 systems. Mutagenesis frequency was calculated by dividing the total number of modified reads by the total number of reads. (A) Comparison of mutagenesis frequency mediated by the Csy4 (blue) and tRNA (beige) based gRNA processing system. The gRNA sites, *Drm1a* gRNA 1 and *Drm1b* gRNAs 1 and 2 were targeted and analyzed by NGS. (B) Comparison of mutagenesis frequency induced by Cas9 (blue) and Cas9_Trex2 (beige). The gRNA sites, *Ms26* gRNA 1, *Ms26* gRNA 2, *Ms45* gRNA1 and *Ms45* gRNA 2, were targeted and analyzed by NGS. (C) Comparison of mutation profiles induced by Cas9 and Cas9_Trex2. The stacked bar graph was generated for each gRNA targeted site with either Cas9 or Cas9_Trex2. The deletions were represented on a grayscale according to size, insertions were represented in red and all other reads (i.e. substitutions, substitutions plus deletions, substitutions plus insertions) were represented in blue. (D) Comparison of DNA repair outcomes induced by Cas9 and Cas9_Trex2. The frequencies of distinct DNA repair outcomes as either MMEJ (blue) or NHEJ (gray) were plotted in each sample treated with Cas9 or Cas9_Trex2.

### The *Trex2* exonuclease enhances targeted mutagenesis with unique mutation profiles

We chose the tRNA-based system for multiplexed genome editing in *S. viridis* for further development due to its proven efficiency and simplicity (Xie, Minkenberg and Yang, 2015; Minkenberg, Wheatley and Yang, 2017). The multiplexing gene editing constructs, pMG198 and pMG199 (Figure S3B), were made containing the Cas9 expression cassette and the gRNA array that simultaneously target two genes, *Ms26* and *Ms45.* When these constructs were tested in protoplasts, high indel mutation frequencies were observed at each target site estimated by Next Generation Sequencing (NGS), i.e. 45% to 60% for the *Ms26* gRNA1 and gRNA2 sites and 35% to 37% for the *Ms45* gRNA1 and gRNA2 sites, respectively (Figure 2B). To test whether the mutagenesis efficiency can be further improved through co-expression of the *Trex2* exonuclease, these multiplexing CRISPR/Cas9 constructs were modified by cloning the *Trex2* coding sequences into the Cas9 expression cassette. The resulting Cas9_Trex2 plasmids, pMG201 and pMG202 (Figure S3B), were then transformed into protoplasts respectively. At each target site, an average of 1.4-fold increase in mutagenesis frequency, ranging from 1.1 to 1.7 folds, was observed from the samples with *Trex2* as compared to those without *Trex2* (Figure 2B). Thus, our results demonstrated that co-expression of the *Trex2* exonucleases with CRISPR/Cas9 further improved mutagenesis frequency in *S. viridis.*

Increased deletion size was observed in tomato and barley when *Trex2* was employed (Čermák *et al*., 2017). However, a thorough characterization of the mutations induced by the combination of Cas9 with the *Trex2* exonuclease has yet to be reported using a large data set. To this end, the mutation profiles were analyzed from a total of 516,815 NGS reads, and compared between the samples with and without co-expression of *Trex2* (Table S2). In the samples without *Trex2,* both insertional and deletional mutations were observed in all 4 targeted sites with 1.6% to 42.1% insertions, the majority of which were 1bp insertions, and 57.1% to 97.8% deletions (Figure 2C). Among these deletions, 97.2% to 98.9%, were smaller than 10 bp. Conversely, in the samples with *Trex2,* essentially no insertions were detected at any target site. On average, 94% of deletions, ranging from 92.3% to 96.6%, were larger than 10 bp with 12% of them extending over 100 bp (Figure 2C). When they were further plotted along each targeted region, the deletions from the samples without *Trex2* were found clustered in 5’ of the PAM sequences (PAM-distal regions) and within 10bp of the DSB site. In contrast, the sequences from the 4 targeted sites with *Trex2* contained much larger deletions that were symmetrically distributed on each side of PAM and that extended up to more than 100 bp (Figure S4).

Interestingly, in the samples with *Trex2,* some specific deletions appeared frequently, exemplified by the 48bp deletions (3.5% of all deletions) in the *Ms26* gRNA2 sample and the 87bp deletions (7.4% of all deletions) in *Ms45* gRNA 2 (Figure S5). Examination of these particular deletions uncovered 2, 3 and 4 bp microhomologies at the *Ms26* gRNA 2 junction sites (Figure S6) and 2, 4, 5 and 6 bp microhomologies at *Ms45* gRNA 2 junction sites, indicating that the microhomology-mediated end joining pathway was involved in creating these deletions. Although previous studies have reported that Cas9-induced DSBs can be repaired through both NHEJ and MMEJ pathway, a recent study indicated that co-expression of *Trex2* with CRISPR/Cas9 predominately results in DSB repair via the NHEJ pathway in human cell lines (Bae *et al*., 2014; Ata *et al*., 2018; Taheri-Ghahfarokhi *et al*., 2018), (Allen *et al*., 2018). To investigate the contribution of these two major end joining pathways in our samples, over 150,000 NGS reads from the samples with and without co-expression of *Trex2* were analyzed based on the presence/absence of microhomology at the deletion junction sites. As a result, in the samples with *Trex2,* a significant fraction of deletions, with an average of 52.2% (ranging from 39.9% to 68.4%) appeared to be repaired by MMEJ, while the samples without *Trex2* exhibited an average of only 3.5% of deletions (ranging from 0.11% to 8.1%) repaired through MMEJ (Figure 2D). These data suggested that different organisms may invoke different end joining pathways to repair the Cas9_Trex2 induced DSBs. Further investigation is needed to better understand the mechanisms underlying these observations and the factors influencing the MMEJ efficiency. Nevertheless, our results indicated that co-expression of the *Trex2* exonuclease with CRISPR/Cas9 can be used as a general strategy to increase the efficiency of targeted deletions, and to create a large collection of mutation variants in plants. Moreover, at least in *S. viridis,* the high frequency of the MMEJ events may also increase the predictability of the mutation outcomes, which is of particular value when precise deletional mutations are needed for dissecting gene function or characterizing cis-regulatory element (Rodríguez-Leal *et al*., 2017; Allen *et al*., 2018).

### Highly efficient multiplexed genome editing in T0 transgenic plants

The multiplex CRISPR/Cas9_Trex2 system tested through the protoplast assay was then used to create heritable mutations in the two linked *Drm1* genes. Three T-DNA constructs were made by assembling the tRNA-gRNA array cassette with the Cas9_Trex2 cassette through the Golden Gate assembly method (Čermák *et al*., 2017). In these T-DNA vectors, the tRNA-gRNA array contained up to 3 gRNAs targeting the *Drm1a* and *Drm1b* genes individually or collectively (Figure S7A). Stable transgenesis was carried out using agrobacterium-mediated transformation. A total of 85, 26, and 103 potential transgenic plants were regenerated from 86, 29 and 112 mature seed-derived calli in the transformation groups, pTW037, pTW044 and pTW045, respectively, exhibiting the high transformation efficiency (above 90%) for this *S. viridis* ME034v variety (Table 1). A subset of candidate plants was randomly picked from each group and further genotyped by genomic PCR using the primers for the hygromycin marker gene (Figure S7B). The presence of T-DNA constructs was confirmed in all 30 plants, indicating a very low escape rate in plant transformation. Notably, the overall transformation efficiency reported in this study has been significantly higher than those from previous studies, i.e. 5-15% for the variety A10.1 (Van Eck, 2018; Nguyen *et al*., 2020). The difference between the observed transformation efficiencies could be attributed to the *S. viridis* variety, ME034v, used in this study. Similarly, high transformation efficiency for ME034v was also observed from other groups (Joyce Van Eck, personal communication). Further investigation will be required to understand the underlying mechanism(s).

**Table 1.** Summary of T0 plant characterization.

| Characterization of T0 plants |  |  |  |  |  |  |
| --- | --- | --- | --- | --- | --- | --- |
| T-DNA Construct | # Calli | # Plants Regenerated | # Plants Genotyped | # Plants with Mutations (Frequency) |  |  |
|  |  |  |  | Drm1a (gRNA1) | Drm1b (gRNA1) | Drm1b (gRNA2) |
| pTW037 | 86 | 85 | 8 (100%)† | NA | 8 (100%) | NA |
| pTW044 | 29 | 26 | 11 (100%)† | 9 (82%) | NA | 8 (73%) |
| pTW045 | 112 | 103 | 11 (100%)† | 11 (100%) | 11 (100%) | 11 (100%) |

† Frequency of transgenic plants indicated in parentheses was confirmed by PCR

Next, the transgenic plants were examined at the targeted regions using genomic PCR followed by restriction enzyme digestion, i.e. the CAPS assay (Figure 3A; Figure S8). As summarized in Table 1, the frequency of plants with mutations induced by the single gRNA T-DNA construct, pTW037, was 100% in the *Drm1a* target site. Similarly, 82% and 73% of plants with the double gRNA T-DNA plasmid, pTW044, carried mutations in the *Drm1a* and *Drm1b* genes; and 100% of plants with the triple gRNA T-DNA construct, pTW045, had mutations in all 3 target sites (Figure 3A and Table 1). Together, the multiplex CRISPR/Cas9_Trex2 system that was demonstrated to be highly efficient in the protoplast assay also induced high frequency mutagenesis in stable transgenic plants.

**Figure 3.**
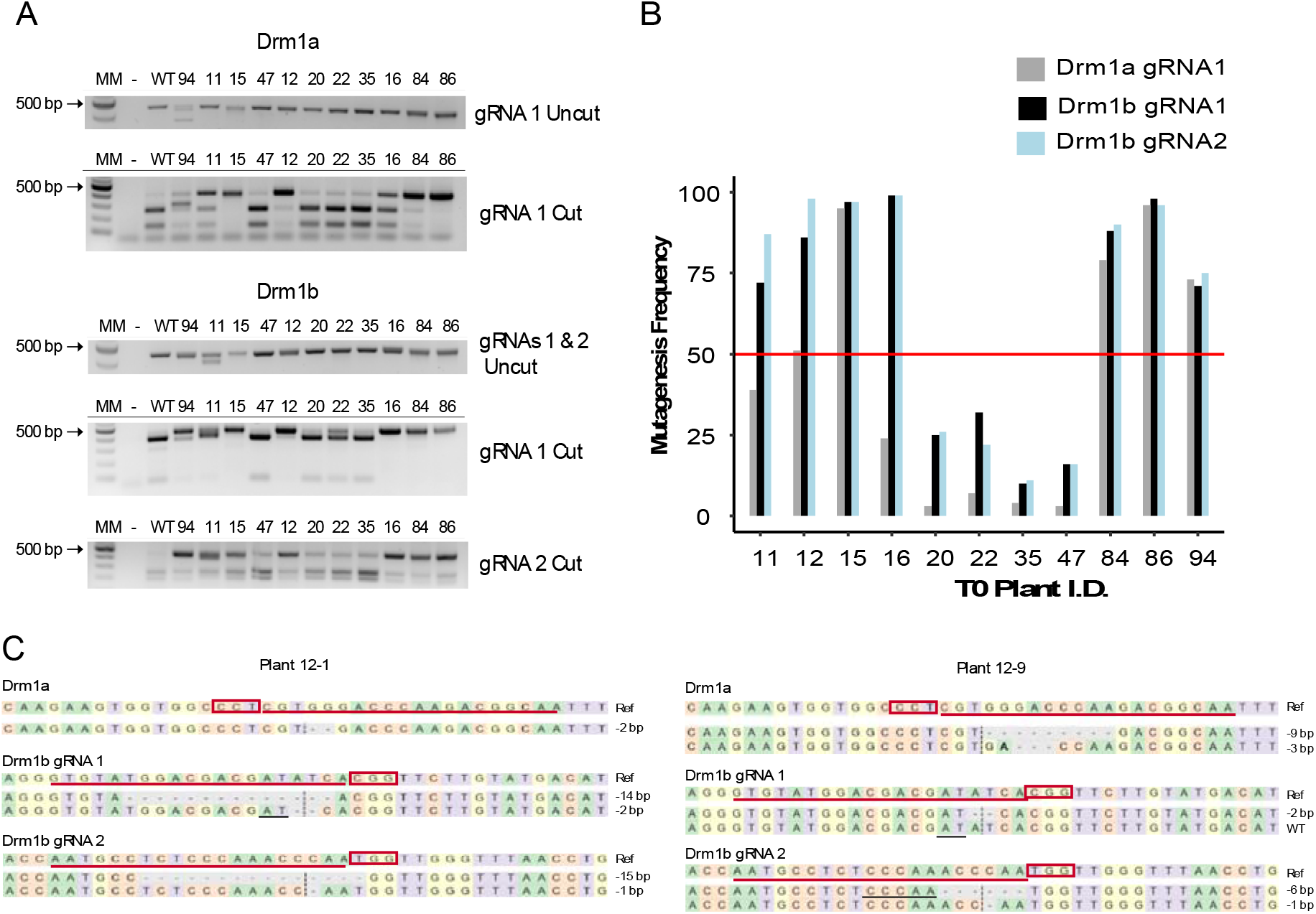
Characterization of mutations induced by the multiplex CRISPR/Cas9_Trex2 system in TO and T1 plants. (A) Genomic PCR and CAPS genotyping assay. Eleven plants (numbered lanes) were genotyped by genomic PCR and the CAPS assay, with a 1kb ladder, no genomic DNA control (-), and wild type DNA control (WT). The results were presented as the PCR amplicons before restriction enzyme digestion (labeled as uncut), and after the digestion (labeled as cut) for each gRNA targeted site. (B) Mutagenesis frequency was determined for each T0 plant by NGS. The mutagenesis frequency was calculated by dividing the total number of modified reads by the total number of reads. The threshold of mutagenesis frequency with 50% was highlighted by the red line. (C) Genotypes of transgene-free T1 plants across all targeted sites. Sequence alignment was shown between the wild type reference sequences, indicated as Ref, and the mutant sequences from Plant 12-1 and 12-9. The gRNA targeted sites were highlighted by the red line with the PAM sequences indicated in the red boxes and the cleavage sites indicated by the vertical dotted lines. All mutations found in these T1 plants were simple deletions. The deleted sequences were indicated by the dashed lines with the size of deletions indicated on the right. The microhomology sequences were highlighted by the black lines.

### Inheritance of targeted mutations in T1 progenies

Plants with the triple gRNA T-DNA construct were further investigated to test heritability of the mutations induced in T0 plants. The CAPS genotyping assay indicated that all eleven T0 plants contained mutations in all three targeted sites at variable frequencies (Figure 3A). To quantify the mutagenesis frequency in each T0 plant, PCR amplicons spanning each targeted region were sequenced using Next Generation Sequencing. Over 20,000 sequencing reads were generated from each PCR amplicon, and analyzed to estimate the indel mutation frequency (Figure 3B). Consistent with results from the CAPS genotyping assay, mutagenesis frequencies were observed in each T0 plant ranging from 3% to 99% in the *Drm1a* site, 10% to 99% in the *Drm1b* gRNA1 site and 11% to 99% in the *Drm1b* gRNA2 site, respectively. In general, the mutagenesis frequencies were positively correlated across all three target sites. For example, four out of eleven plants showing lower mutagenesis efficiency in the *Drm1a* target site (under 10%) also displayed lower mutagenesis efficiency in the two *Drm1b* target sites (Figure 3B).

Five T0 plants, 12,15, 84, 86, and 94, showing mutation frequencies greater than 50% at all three target sites, were chosen to be self-pollinated and grown to maturity. Ten T1 progenies from each T0 plant were grown for further characterization. Using the CAPS genotyping assay, high frequencies of mutant plants were detected in these T1 populations, ranging from 50-100%, 60-90%, and 30-100% at three target sites, *Drm1a, Drm1b* gRNA1 and *Drm1b* gRNA2, respectively (Figure S9, Table S3). To further distinguish heritable mutations from somatic mutations in these T1 plants, genomic PCR was conducted to detect T-DNA transgene free plants using primers for the hygromycin marker gene. Four out of five T1 populations, 12,15, 84 and 86, exhibited segregation of the transgene (Table 2, Figure S10). Out of 50 T1 plants, 10 transgene-free plants were identified (Table 3). Among them, 6 plants (60%) showed mutations at at least one of the three target sites, with 1 plant (10%) having a mutation at two target sites and 2 plants (20%) having mutations at all three target sites (Table 3). These two transgene free plants, 12-1 and 12-9, with mutations in all three target sites were further characterized by NGS (Table 3, Figure 3C). Notably, while the CAPS assay suggested that heterozygous mutations occurred at the *Drm1b* gRNA2 site in these plants (Figure S9), the NGS data revealed bi-allelic mutations in both plants. Close examination of these mutations identified a 1bp deletion at this target site that did not disrupt the restriction enzyme (*PfIM*I) recognition site used in the CAPS assay. This finding suggested that the CAPS assay used to screen the T1 plants may underestimate the mutagenesis frequency transmitted to the T1 populations. Additionally, as seen from the protoplast assay, MMEJ-mediated deletions were recovered in each plant (Figure 3C). Taken together, these results clearly demonstrated that the mutations are transmissible in the *S. viridis* mutant plants at high efficiency. This made it possible to recover homozygous mutants from a relatively small population of T1 plants and to generate multiple gene knock-out events simultaneously and rapidly, which is particularly useful in editing tightly linked genetic loci as shown in this study.

**Table 2.** Summary of T1 plant characterization.

| Characterization of T1 plants |  |  |  |  |  |
| --- | --- | --- | --- | --- | --- |
| T0 Plant I.D. | # T1 Plants | # Transgene-free T1 | Number of Plants with mutations (Frequency) |  |  |
|  |  |  | Drm1a (gRNA1) | Drm1b (gRNA1) | Drm1b (gRNA2) |
| 12 | 10 | 5 | 5 (50%) | 6 (60%) | 3 (30%) |
| 15 | 10 | 2 | 8 (80%) | 8 (80%) | 9 (90%) |
| 84 | 10 | 2 | 8 (80%) | 8 (80%) | 9 (90%) |
| 86 | 10 | 1 | 10 (100%) | 9 (90%) | 10 (100%) |
| 94 | 10 | 0 | 7 (70%) | 8 (80%) | 10 (100%) |

**Table 3.** Summary of transgene-free T1 plant characterization.

| Genotype Characterization of Transgene-free T1 Plants |  |  |  |
| --- | --- | --- | --- |
| Plant ID | *Drm1a* | *Drm1b* target 1 | *Drm1b* target 2 |
| 12-1 | homozygous (-2bp/-2bp)† | bi-allelic (-14bp/-2bp)† | bi-allelic (-15bp/-1bp)† |
| 12-9 | bi-allelic (-9bp/-3bp)† | heterozygous (-2bp/WT)† | bi-allelic (-6bp/-1bp)† |
| 12-15 | WT‡ | WT | WT |
| 12-17 | WT | WT | WT |
| 12-22 | WT | WT | WT |
| 15-88 | WT | WT | heterozygous |
| 15-95 | WT | WT | heterozygous |
| 84-18 | WT | WT | heterozygous |
| 84-27 | WT | WT | WT |
| 86-13 | heterozygous | WT | heterozygous |

† Genotypes were confirmed by Next Generation Sequencing. The size of deletion was indicated in the parentheses.
‡ WT meant the wild type sequence without mutations.

In summary, we developed a protoplast-based assay to rapidly test and optimize the multiplex CRISPR/Cas9 gene editing system in *S. viridis.* The mutagenesis frequency can be enhanced by co-expression with the *Trex2* exonuclease resulting in a unique mutation profile with larger deletions, no insertions and a high frequency of MMEJ repaired events. Further, the optimized multiplex CRISPR/Cas9_Trex2 system can induce targeted mutagenesis in stable transgenic plants at remarkably high efficiency. This system allowed us to generate heritable knock-outs in two tightly-linked *S. viridis* genes from a small number of transgenic plants (10 T0 plants) in a timeframe as short as three months (starting from plant transformation to T1 seedlings). With the efficient agrobacterium-mediated transformation method, this highly efficient pipeline makes it possible to create a large mutant collection of *S. viridis* in a rapid and high throughput manner. Moving forward, it would be interesting to test this CRISPR/Cas9_Trex2 system, or combinations of *Trex2* with different Cas proteins, in other plants. These new systems have the potential to be widely applicable in achieving more predictable MMEJ-mediated mutations in many plant species.

## Experimental procedures

### Plant materials, seed germination and plant growth conditions

S. *viridis* variety ME034v was used in this study. To break dormancy and promote seedling germination, freshly harvested seeds were incubated at 29□c for 24 hours in a 1.4 mM gibberellic acid and 30 mM potassium nitrate solution (Sebastian *et al*., 2014). After the 24-hour incubation, seeds were sterilized with 50% bleach for 10 minutes, followed by 5 water rinses and planted on germination media (0.5X MS, 0.5% sucrose, 0.4% PhytaGel, pH 5.7). Seedlings were transplanted to soil six days after germination and grown at 26°C/22°C (day/night) with a photoperiod of 16h/8h (day/night), under 30% relative humidity, a modified protocol from (Huang *et al*., 2019).

### Guide RNA design and vector construction

The genomic sequences of each targeted gene were obtained by BLAST searching the *S. viridis* A10.1 reference genome from the phytozome database (https://phytozome.jgi.doe.gov). CRISPR gRNAs were designed to target exons in the 5’ region of the gene or the conserved domains in each gene using CRISPOR (Haeussler *et al*., 2016). The targeted sequences were further verified by Sanger sequencing in the *S. viridis* variety ME034v. The conserved domains were identified by comparing the coding sequences from *S. viridis* with their orthologs from brachypodium, maize, and Arabidopsis.

The gRNA constructs were made by following the Golden Gate assembly method (Čermák *et al*., 2017). The backbone for the tRNA-based gRNA construct was pMOD_B2103, and the backbone of the Csy4-based gRNA construct was pMOD_B2303. The Cas9 constructs were pMOD_A1110 and pMOD_A1510. The Cas9_Trex2 construct, pMOD_A1910, were made by cloning the Trex2 coding sequence into the codon-optimized Cas9 expression cassette, based on the codon usage from wheat (*Triticum aestivum*). The GFP reporter constructs, pMOD_C3003 and pMOD_C3013, were made by cloning the GFP coding sequences under the control of the *CmYLCV* and *PvUbi* promoters with the 35S terminator. All the constructs will be deposited to Addgene.

### Protoplast isolation and transformation

Protoplast isolation and transformation were performed using a modified version of the PEG-mediated method (Li *et al*., 2016). In brief, leaves from 14-day young seedlings were sliced into small pieces with a razor blade and digested with the enzyme solution (1.5% Cellulase, 0.75% Macerozyme, Kanematsu USA Inc.) for 4-5 hours on a shaker at 40 rpm. The digested tissues were filtered through a 70uM nylon filter (Fisher Scientific LLC) into W5 buffer (2mM MES with pH5.7, 154mM NaCl, 125mM CaCI_2_, 5mM KCl). Protoplasts were collected and resuspended in W5 buffer with a gentle centrifuge at 100xg for 5 minutes. The number of protoplasts was estimated using a hemocytometer. Roughly 200,000 protoplasts were mixed with DNA plasmids (15ug per construct) in 20% PEG buffer and incubated at room temperature in the dark for 48 hours. Transformation efficiencies were monitored by transforming protoplasts with a plasmid encoding GFP.

### T-DNA transformation and tissue culture

*Agrobacterium tumefaciens-mediated* transformation of *S. viridis* was performed as previously described with a few modifications (Van Eck *et al*., 2017). Callus initiation was first performed by removing the seed coats and sterilizing seeds with a 10% bleach plus 0.1% tween solution for 5-10 minutes under gentle agitation. Seeds were plated on callus induction media with the embryos facing upward. The plates were placed at 24°C in the light for a week and then moved to dark for callus initiation. Embryogenic calli were collected after 4-7 weeks and inoculated with the AGL1 strain harboring the T-DNA construct. Inoculated calli were placed on co-culture medium and incubated in the dark at 20°C for 5-7 days. Transformed calli were transferred to the selection medium with 50mg hygromycin for 4 weeks at 24°C, then the selected calli were subcultured on plant regeneration media with 20mg hygromycin under 16-hour light to allow the growth of the transformed shoots. Elongated shoots were transferred to the rooting medium with 20mg hygromycin. Shoots with well-developed roots were transplanted to soil and grown to maturity.

### Genotyping and mutant identification

Mutant identification and characterization were performed using two methods, genomic PCR with restriction enzyme digestion (CAPS assay) and Illumina paired-end read amplicon sequencing (NGS assay). PCR was performed with GoTaq Green Master Mix (Promega Inc.) according to the manufacturer’s instructions with an annealing temperature of 58□C and an extension time of one minute. Amplicons were then subjected to restriction enzyme digestion using an enzyme that overlaps with the CRISPR/Cas9 cleavage site. PCR amplicons made with the corresponding primers were subjected to Illumina paired-end read amplicon sequencing by Genewiz Inc. The raw NGS reads were analyzed using CRISPResso2 (Clement *et al*., 2019). All the primers used in this study were listed in Table S4.

### Characterization of mutation profiles

Mutations in the NGS reads were characterized into three categories, deletions, insertions and others (including substitutions and substitutions with insertions or deletions). The mutagenesis efficiency of each mutation type was estimated by dividing the total number of modified reads by the total number of reads. To minimize the problem caused by sequencing errors from NGS, mutation reads that only occurred once in the NGS data were not included in the calculation. To quantify the mutations derived from NHEJ or MMEJ repair pathways, each distinct deletion read was categorized into three separate sequences: 1. the left flanking sequence, 2. the deleted sequence, and 3. the right flanking sequence. Mutation reads were considered as MMEJ products when greater than 2 bp of homology were identified at the junction site between left and right flanking sequences. Mutation reads without microhomology sequences at the junction sites were classified as NHEJ events.

## Acknowledgments

We thank all members of Voytas Lab and Zhang Lab for helpful discussions and feedback, and Meredith Song for assisting with genomic DNA extractions, genotyping and caring for plants. F.Z. was supported by the startup fund to F.Z from the College of Biological Sciences, University of Minnesota. D.V, C.S and M.G were supported in part by a grant DE-SC0018277 from the DOE Department of Biological and Environmental Research. H.Z was supported by the grant ZDYF2017017 from the key research and development project of Hainan. P.Z., P.C. and N.M.S. were supported in part by a grant from NSF (IOS-1802848). University of Minnesota Department of Plant and Microbial Biology and Department of Genetics, Cell Biology and Development are part of a team supporting DARPA’s LiSTENS Program.

## Conflicts of interest

The authors declare no conflict of interest.

## Supplemental Figure Legends

**Figure S1.**
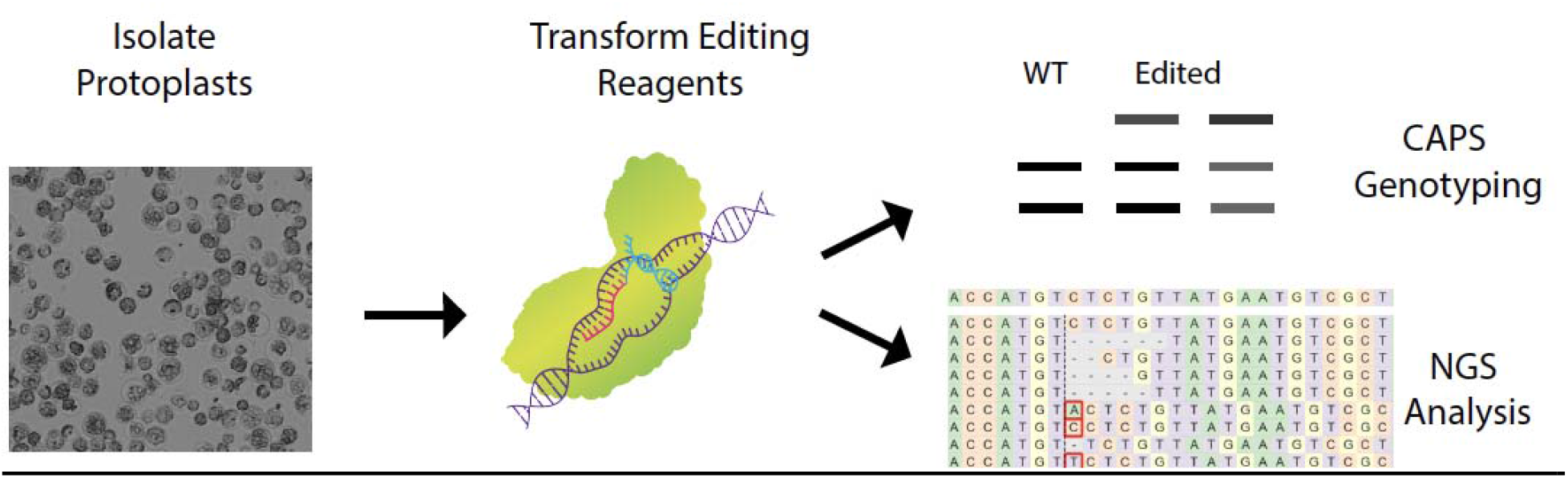
*Setaria viridis* protoplast assay pipeline to test genome editing reagents. Following protoplast isolation from young leaves of 14-day old plants, genome editing reagents can then be transformed into protoplasts to evaluate editing efficacy. At 48 hpt, genomic DNA is extracted, subjected to Cleaved Amplified Polymorphic Sequences (CAPS) or Next Generation Sequencing (NGS) analysis.

**Figure S2.**
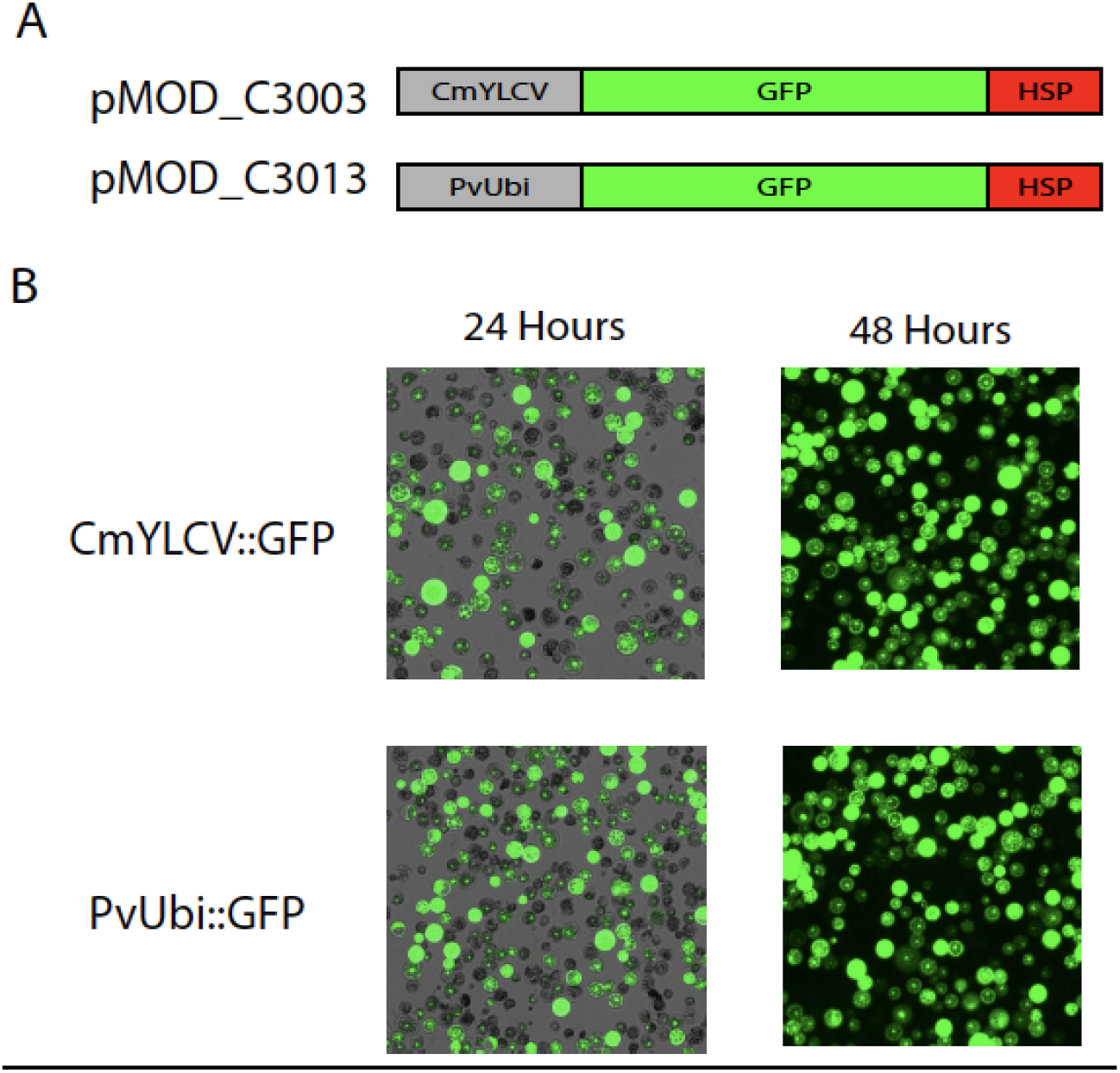
(A) The schematic structure of the GFP reporter plasmids used in the protoplast. The GFP coding sequence (green boxes) was driven by either the *CmYLCV* (plasmid ID pMOD_C3003) or the *PvUbi2* promoter (plasmid ID pMOD_C3103) labeled as the gray boxes with the HSP terminator (red boxes). The illustration was not to scale. (B) Mesophyll protoplast cells isolated from young leaves and transformed with the GFP reporter plasmids, pMOD_C3003 and pMOD_C3013. GFP expression was assayed at 24 and 48 hpt.

**Figure S3.**
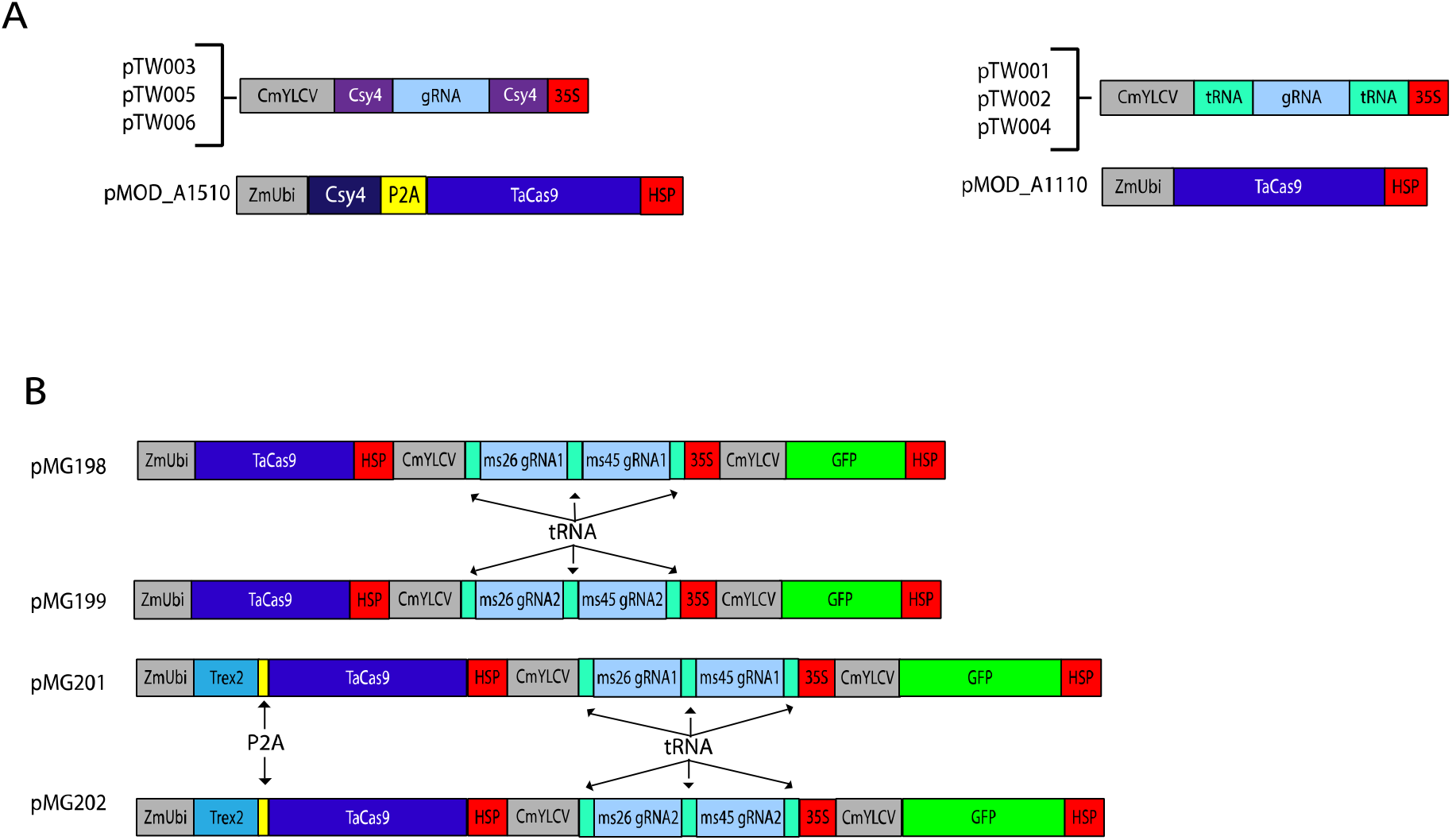
(A) The schematic structures of the plasmids to test the Csy and tRNA-based gRNA processing systems. In the Csy4 system, the Cas9 expressing plasmid, pMOD_A1510, contained the Cys4 coding sequence (dark blue box) with the Cas9 coding sequence (blue box) separated by the P2A sequence (yellow box) under the *ZmUbi* promoter (grey box). The gRNA expressing plasmids, pTW003, pTW005and pTW006, contained a single gRNA sequence (light blue box) flanked by the Csy4 recognition sites (purple boxes) under the control of the *CmYLCV* promoter and the 35S terminator (red box). In the tRNA system, the Cas9 expressing plasmid, pMOD_A1110, contained only the Cas9 coding sequences driven by the *ZmUbi* promoter. The gRNA expression plasmids, pTW001, pTW002 and pTW004, contained a single gRNA sequence (light blue box) flanked by the tRNA sequences (light green boxes). (B) The schematic structures of the plasmids to the multiplexed Cas9 and Cas9_Trex2 systems. The plasmids, pMG198, pMG199, pMG201 and pMG202, were constructed to contain three components, the Cas9 or Cas9_Trex2 expression cassette, the tRNA based multiplexing gRNA cassette, and the GFP reporter. The illustration was not to scale.

**Figure S4.**
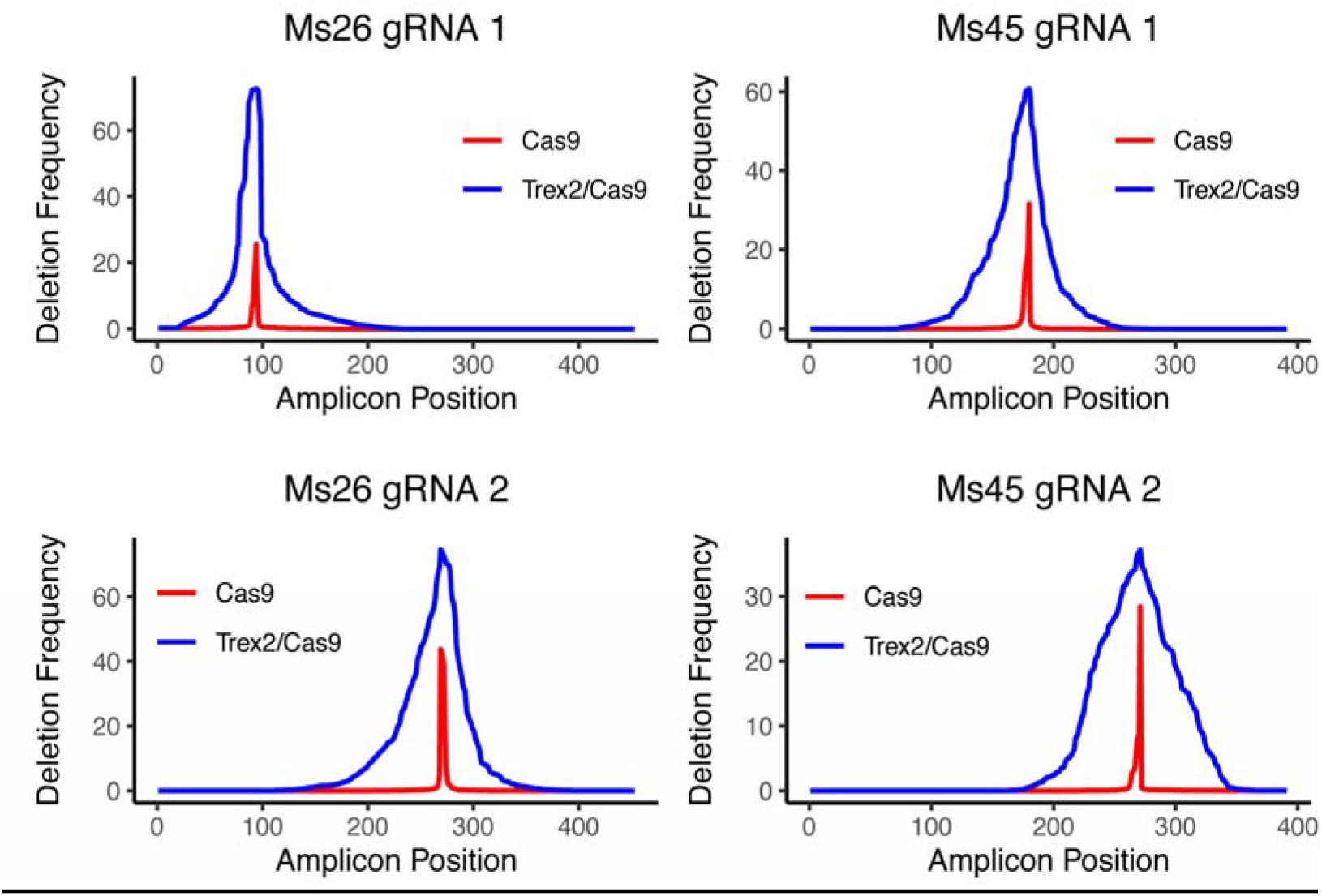
Distribution of deletions across the 400bp - 500bp of the gRNA targeted regions induced by either Cas9 (red) and Cas9_Trex2 (blue). The Deletion frequency was calculated by dividing the total number of deletions at each nucleotide by the total number of deletions reads.

**Figure S5.**
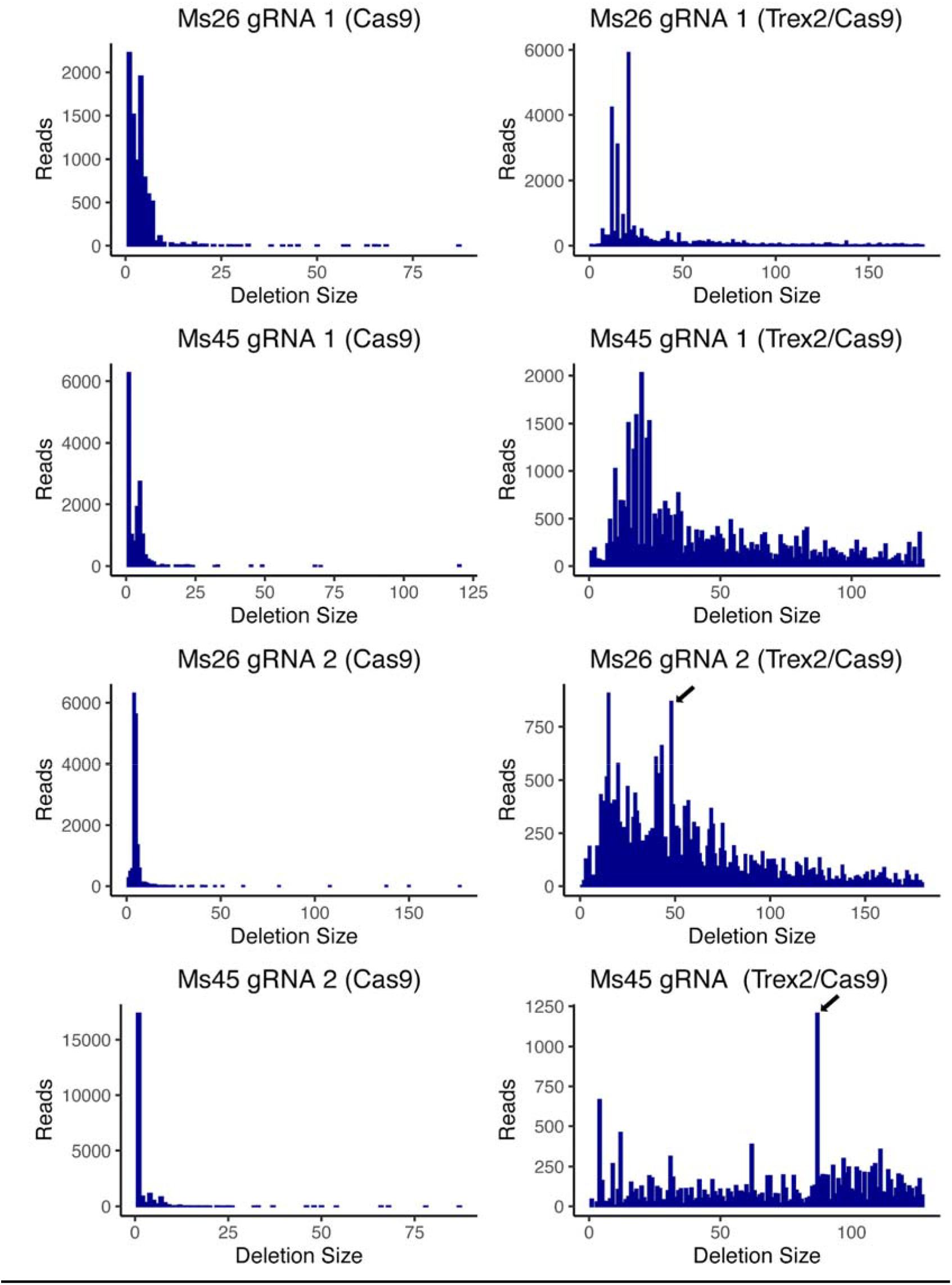
Comparison of the size distribution from deletions induced by Cas9 or Cas9_Trex2. The number of reads for each deletion size was estimated for 4 gRNA sites from the Ms26 and Ms45 genes, respectively. The examples of MMEJ-mediated deletions were indicated by the black arrows.

**Figure S6.**
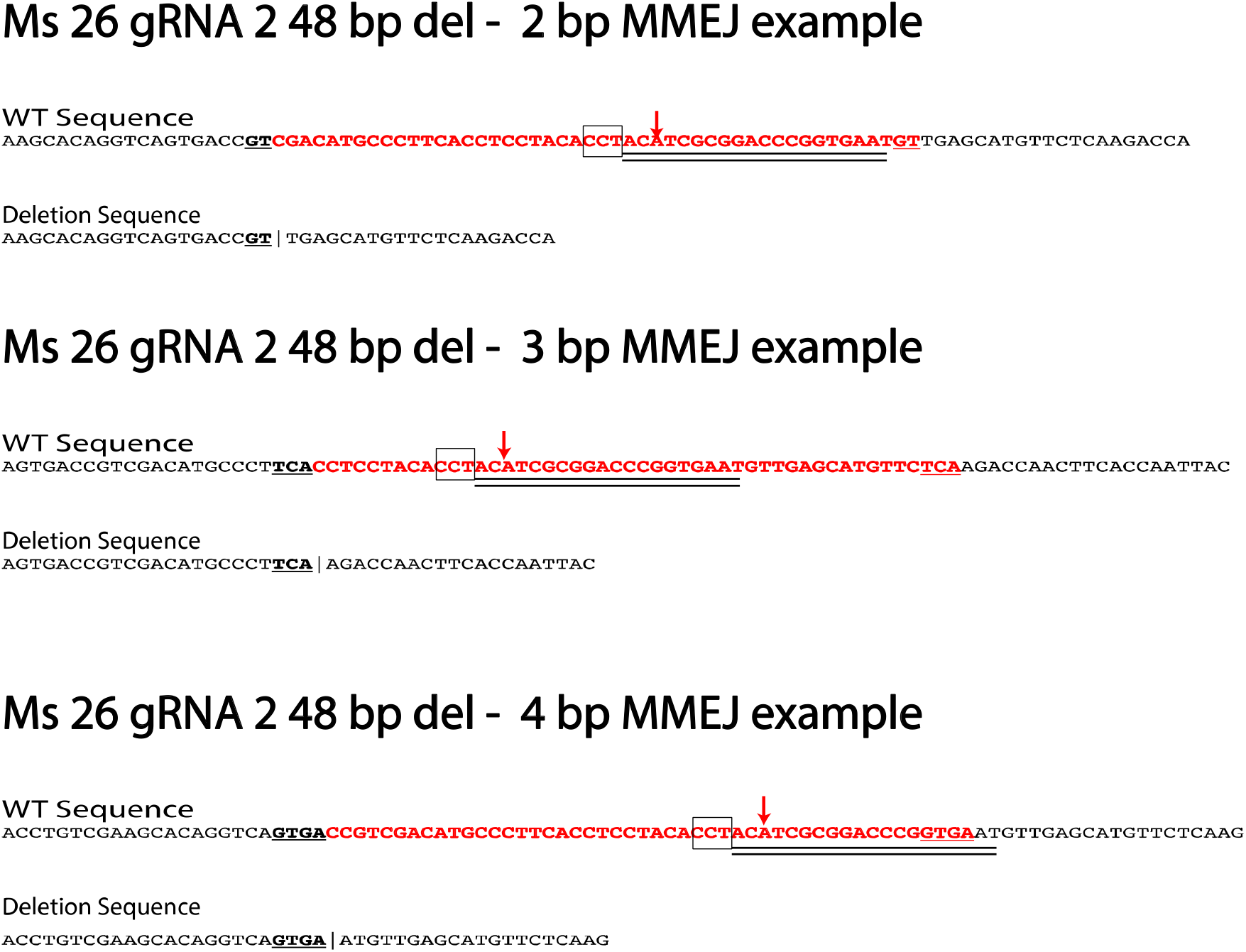
Example of the MMEJ-mediated deletions. The CRISPR gRNA target sites were indicated by the black double lines with the PAM sequences outlined with the black boxes. The CRISPR/Cas9 cleavage sites were pointed by the red arrows. The microhomology sequences were underlined in the wild type sequences with the deleted sequences highlighted in red.

**Figure S7.**
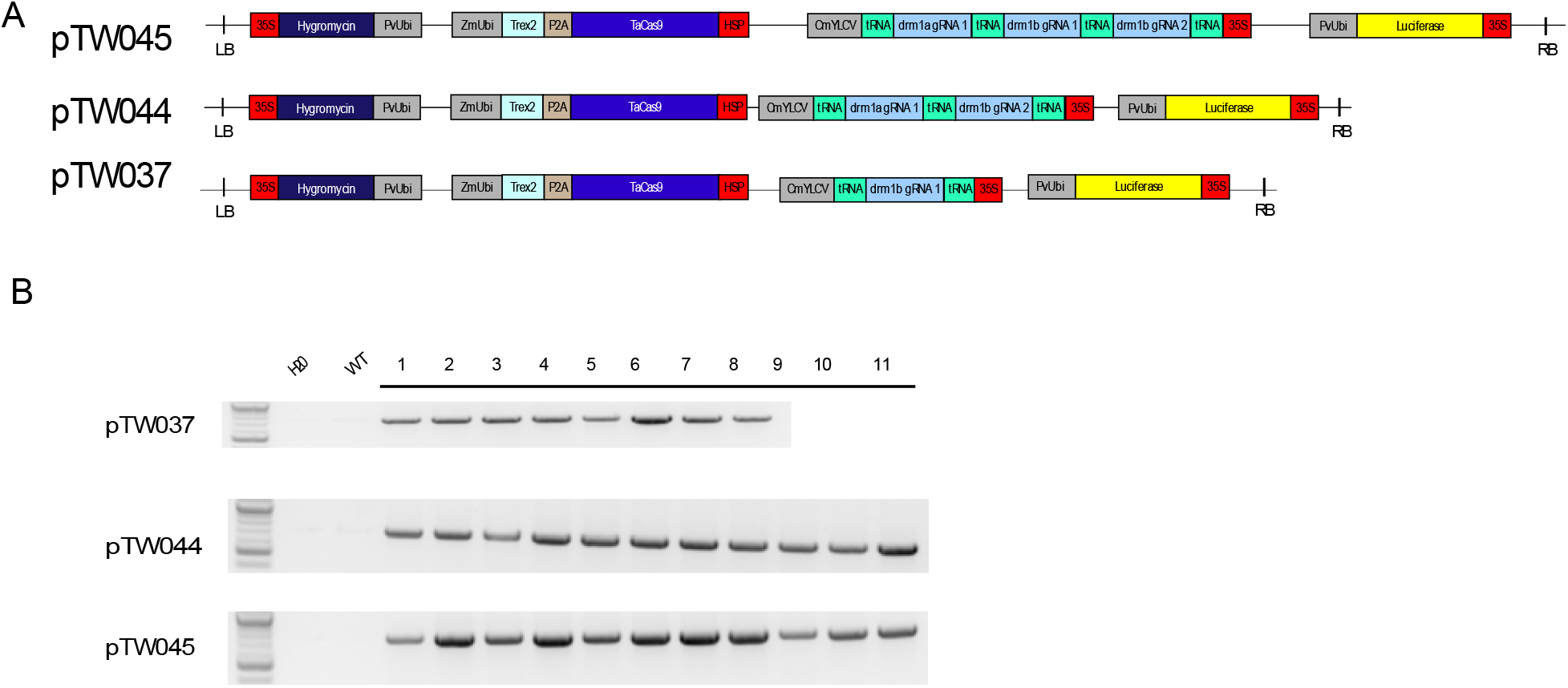
(A) The schematic structures of T-DNA binary plasmids for stable transgenesis. Each T-DNA plasmid contained three components, the Cas9_Trex2 expression cassette, the tRNA-based multiplexing gRNA cassette, and the luciferase reporter. Within each construct, pTW037, contained one gRNA targeting Drm1b, pTW044 contained one gRNA targeting Drm1a and one gRNA targeting Drm1b, and pTW045 contained one gRNA targeting Drm1a and two gRNAs targeting Drm1b. (B) Genomic PCR genotyping to detect the presence of the T-DNA in T0 plants. Two controls were included in this experiment, one without genomic DNA (indicated as H_2_O) and the other with wild type *S. viridis* genomic DNA (indicated as WT).

**Figure S8.**
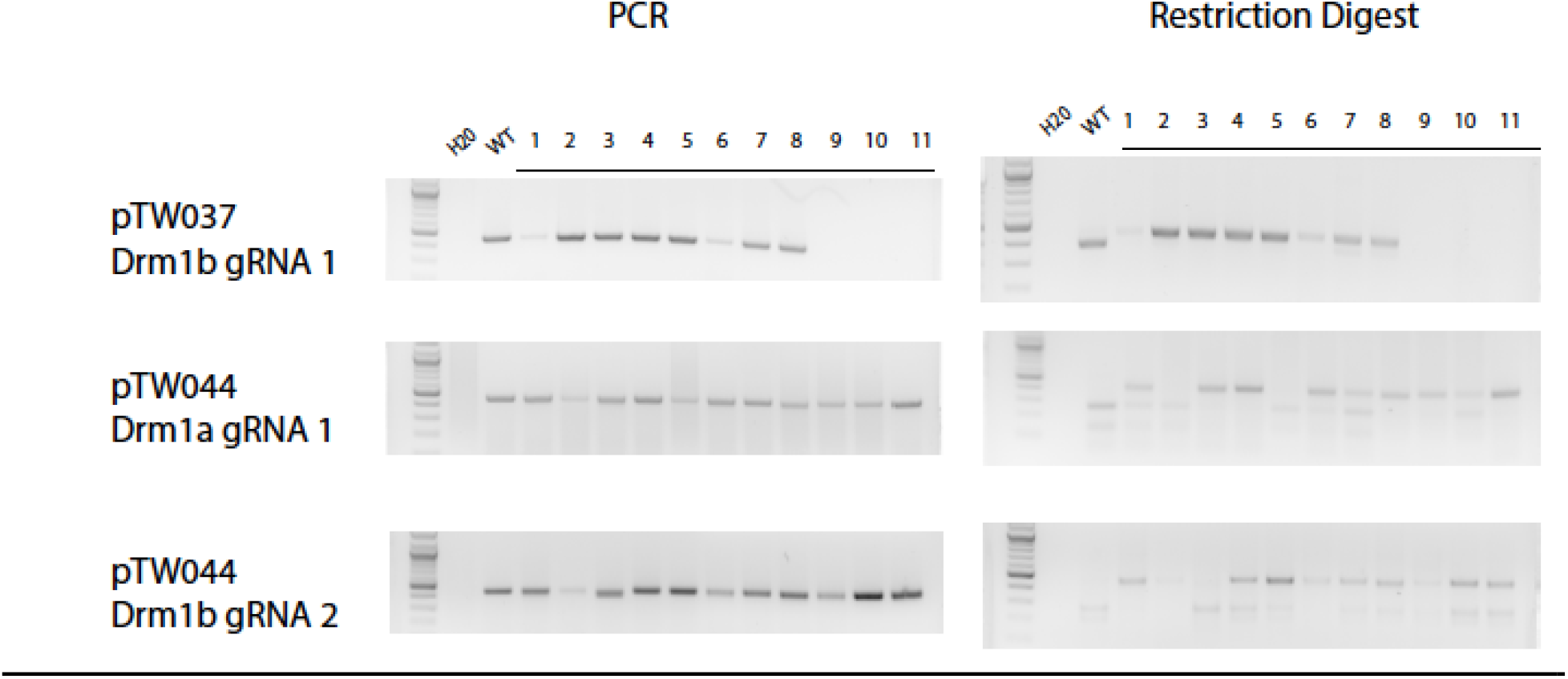
Genomic PCR genotyping of T0 plants with the CAPS assay. The samples from left to right were a 1 kb ladder, a no-genomic DNA control (indicated as H_2_0), a control with wild type genomic DNA (indicated as WT) and individual T0 samples.

**Figure S9.**
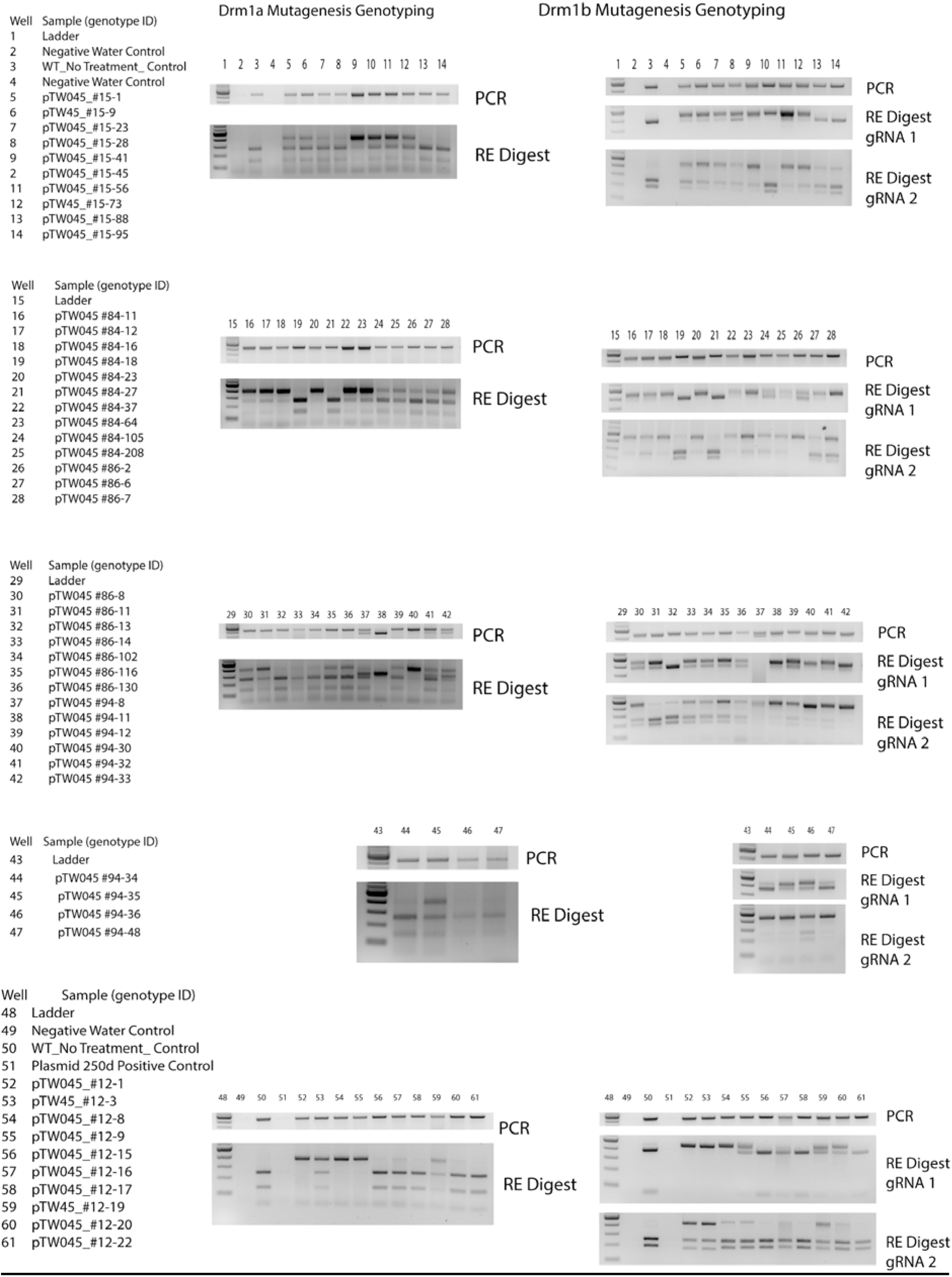
Genomic PCR genotyping of T1 plants with the CAPS assay. The samples from left to right were 1 kb ladder, no-genomic DNA control (H_2_0) and wild type genomic DNA control (WT). The remaining samples corresponded to individual T1 plants.

**Figure S10.**
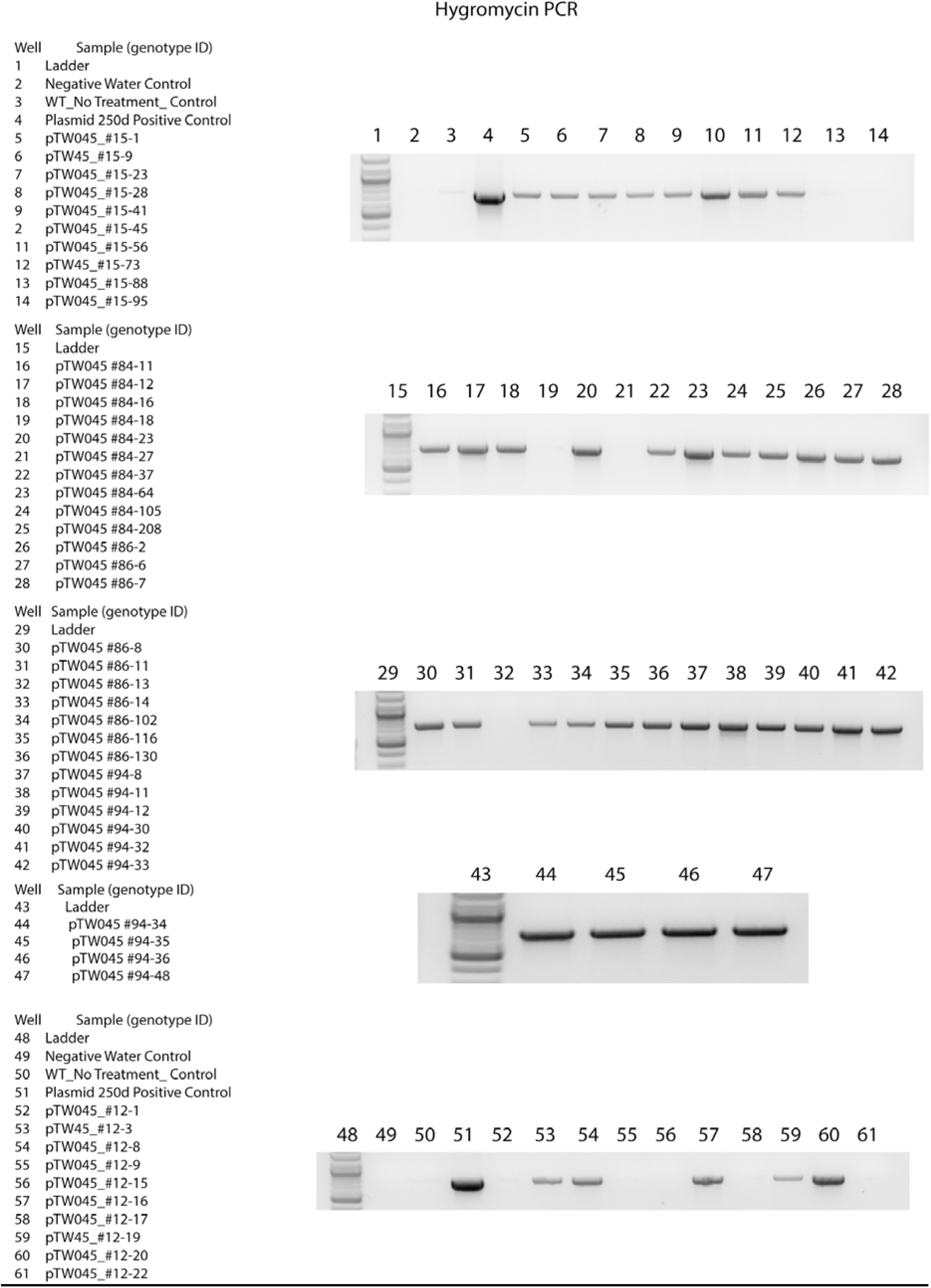
Genomic PCR genotyping for segregation of the T-DNA transgenes in T1 plants. The samples from left to right were 1 kb ladder, no-genomic DNA control (H_2_0), and wild type genomic DNA control (WT) and individual T1 samples.

**Table S1.**
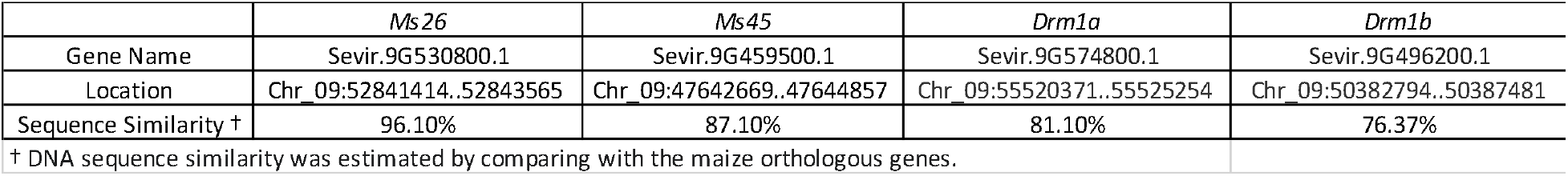
Summary of the targeted genes.

**Table S2.** Summary of Next Generation Sequencing reads for each targeted site.

| Gene (gRNA) | Trex2 Co-expression | Total |  | Unique |  |
| --- | --- | --- | --- | --- | --- |
|  |  | Reads | Modified Reads | Modified Read | Deletions |
| Ms26 (gRNA 1) | No | 32,088 | 25,802 | 991 | 978 |
| Ms26 (gRNA 2) | No | 55,292 | 33,382 | 1,684 | 1,658 |
| Ms45 (gRNA 1) | No | 63,134 | 23,230 | 1,367 | 1,354 |
| Ms45 (gRNA 2) | No | 80,344 | 27,766 | 1,436 | 1,427 |
| Ms26 (gRNA 1) | Yes | 60,676 | 44,793 | 3,766 | 3,751 |
| Ms26 (gRNA 2) | Yes | 60,578 | 45,669 | 3,633 | 3,620 |
| Ms45 (gRNA 1) | Yes | 96,953 | 59,862 | 3,803 | 3,709 |
| Ms45 (gRNA 2) | Yes | 67,750 | 25,872 | 2,027 | 2,023 |

**Table S3.** Summary of T1 plant genotypes.

| Plant ID | Transgenic | T1 Genome Editing Characterization |  |  |
| --- | --- | --- | --- | --- |
|  |  | *Drm1a* | *Drm1b* target 1 | *Drm1b* target 2 |
| 12-1 | no | bi-allelic or homozygous | bi-allelic or homozygous | heterozygous |
| 12-3 | yes | heterozygous | bi-allelic or homozygous | heterozygous |
| 12-8 | yes | bi-allelic or homozygous | bi-allelic or homozygous | no mutation |
| 12-9 | no | bi-allelic or homozygous | heterozygous | no mutation |
| 12-15 | no | no mutation | no mutation | no mutation |
| 12-16 | yes | no mutation | no mutation | no mutation |
| 12-17 | no | no mutation | no mutation | no mutation |
| 12-19 | yes | heterozygous | heterozygous | heterozygous |
| 12-20 | yes | no mutation | heterozygous | no mutation |
| 12-22 | no | no mutation | no mutation | no mutation |
| 15-1 | yes | heterozygous | heterozygous | heterozygous |
| 15-9 | yes | heterozygous | heterozygous | heterozygous |
| 15-23 | yes | heterozygous | heterozygous | heterozygous |
| 15-28 | yes | heterozygous | heterozygous | heterozygous |
| 15-41 | yes | heterozygous | bi-allelic or homozygous | heterozygous |
| 15-45 | yes | heterozygous | bi-allelic or homozygous | no mutation |
| 15-56 | yes | heterozygous | heterozygous | heterozygous |
| 15-73 | yes | heterozygous | heterozygous | heterozygous |
| 15-88 | no | no mutation | no mutation | heterozygous |
| 15-95 | no | no mutation | no mutation | heterozygous |
| 84-11 | yes | heterozygous | heterozygous | heterozygous |
| 84-12 | yes | heterozygous | heterozygous | heterozygous |
| 84-16 | yes | heterozygous | heterozygous | heterozygous |
| 84-18 | no | no mutation | no mutation | heterozygous |
| 84-23 | yes | bi-allelic or homozygous | bi-allelic or homozygous | heterozygous |
| 84-27 | no | no mutation | no mutation | no mutation |
| 84-37 | yes | heterozygous | heterozygous | heterozygous |
| 84-64 | yes | heterozygous | heterozygous | heterozygous |
| 84-105 | yes | heterozygous | heterozygous | heterozygous |
| 84-208 | yes | heterozygous | heterozygous | heterozygous |
| 86-2 | yes | heterozygous | heterozygous | bi-allelic or homozygous |
| 86-6 | yes | heterozygous | heterozygous | heterozygous |
| 86-7 | yes | heterozygous | bi-allelic or homozygous | heterozygous |
| 86-8 | yes | heterozygous | heterozygous | heterozygous |
| 86-11 | yes | heterozygous | heterozygous | heterozygous |
| 86-13 | no | heterozygous | no mutation | heterozygous |
| 86-14 | yes | heterozygous | heterozygous | heterozygous |
| 86-102 | yes | heterozygous | heterozygous | heterozygous |
| 86-116 | yes | heterozygous | heterozygous | heterozygous |
| 86-130 | yes | heterozygous | heterozygous | heterozygous |
| 94-8 | yes | heterozygous | heterozygous | heterozygous |
| 94-11 | yes | bi-allelic or homozygous | no mutation | heterozygous |
| 94-12 | yes | heterozygous | heterozygous | heterozygous |
| 94-30 | yes | bi-allelic or homozygous | no mutation | heterozygous |
| 94-32 | yes | heterozygous | heterozygous | heterozygous |
| 94-33 | yes | heterozygous | heterozygous | bi-allelic or homozygous |
| 94-34 | yes | no mutation | heterozygous | bi-allelic or homozygous |
| 94-35 | yes | heterozygous | heterozygous | bi-allelic or homozygous |
| 94-36 | yes | no mutation | heterozygous | heterozygous |
| 94-48 | yes | no mutation | heterozygous | bi-allelic or homozygous |

**Table S4.**
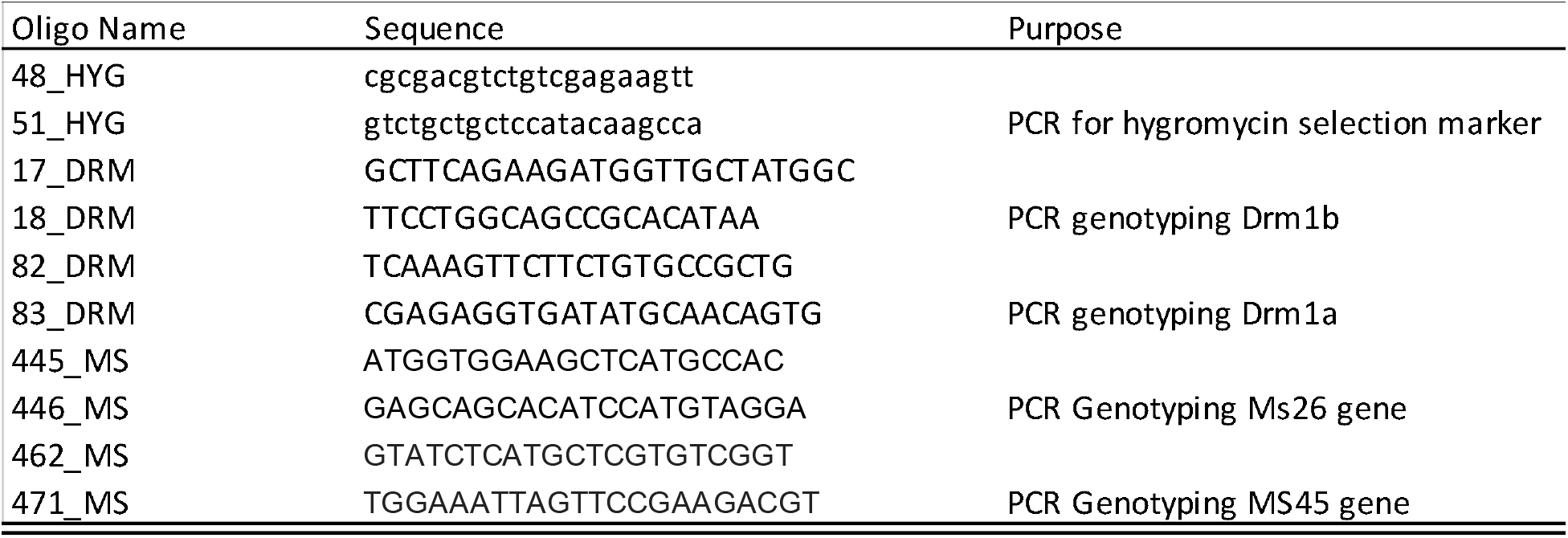
Summary of the primer information.

## Notes

### Competing Interest Statement

The authors have declared no competing interest.

